# Release of HIV-1 particles from the viral compartment in macrophages requires an associated cytoskeleton and is driven by mechanical constraints

**DOI:** 10.1101/2021.12.31.474638

**Authors:** Vasco Rodrigues, Sarah Taheraly, Mathieu Maurin, Mabel San-Roman, Emma Granier, Anaël Hanouna, Phillippe Benaroch

## Abstract

A defining feature of HIV-1 replication in macrophages is that viral assembly occurs at the limiting membrane of a compartment often named VCC (virus-containing compartments) that is connected to the extracellular medium. The newly formed viral progeny pinches of the membrane and accumulates in the lumen of the VCC. While HIV budding has been extensively studied, very little is known about how viral particles present in the lumen of VCC are released in the extracellular medium. Here we show that the actin dynamics are critical for this process by combining ultrastructural analyses, time-lapse microscopy and perturbations of the actin cytoskeleton. We found that jasplakinolide, which stabilizes actin fibres, inhibited viral release from HIV-1-infected macrophages. Furthermore, in jasplakinolide-treated macrophages, VCC became scattered and no longer co-localized with the integrin CD18, nor the phosphorylated form of the focal adhesion kinase PYK2. Inhibition of PYK2 activity in infected macrophages promoted intracellular retention of viral particles in VCC that were no longer connected to the plasma membrane. Finally, we stimulated the rapid release of viral particles from the VCC by subjecting infected macrophages to frustrated phagocytosis. As macrophages spread on IgG-coated glass surfaces, VCC rapidly migrated to the basal membrane and released their viral content in the extracellular medium, which required their association with CD18 and the actin cytoskeleton. These results highlight that VCC trafficking and virus release are intimately linked to the reorganization of the macrophage actin cytoskeleton in response to external physical cues, suggesting that it might be regulated in tissues by the mechanical stress to which these cells are exposed.

## Introduction

Macrophages are embedded in many tissues where they ensure specific functions. Early studies established that HIV-1-infected patients possess infected macrophages in many of their tissues (Orenstein et al., 1988). Usage of monocyte-derived macrophages (MDM) exposed to HIV-1 *in vitro* allowed the study of the viral replication cycle and revealed some of its specific features as compared to how it replicated in T lymphocytes (Rodrigues et al., 2017; Sattentau and Stevenson, 2016). Viral assembly takes place in macrophages at the limiting membrane of a compartment that appears intracellular. The newly formed virions, pinch off from this membrane in the lumen of the compartment, hence called VCC for virus-containing compartments, where they accumulate. Interestingly, compartments with very similar characteristics are also present in uninfected macrophages, as they similarly express tetraspanins (CD9, CD81 and CD53) (Deneka et al., 2007), and the scavenger receptor CD36 (Berre et al., 2013). Upon HIV-1 infection, newly synthesized Gag is recruited to these pre-existing compartments that become *de facto* VCC (Berre et al., 2013).

The VCC possess a very intricate architecture and are often connected to the extracellular medium by channels or conduits that are too narrow to allow the virions to exit but that permit fluid exchanges (Gaudin et al., 2013; Welsch et al., 2007). They remain mostly inaccessible to macromolecules delivered from the extracellular media, such as antibodies (Chu et al., 2012). The precise nature of the VCC has been debated but the current view holds that it originates from tetraspanin-rich zones of the plasma membrane that have been internally sequestered. While VCC are generally considered as continuous with the plasma membrane, recent work also evidenced the presence of completely enclosed compartments that could fuse with plasma membrane-connected compartments (Ladinsky et al., 2019). These studies highlight a rather complex and dynamic structure that results in a more complex release of viral particles as compared with other cell types.

How the virions stocked in the VCC lumen are released in the extracellular medium remains an open question. To our knowledge, a single study reported that the release of viral particles from the VCC can be induced. Exposure of an infected MDM to extracellular ATP (eATP) prompted a rapid discharge of the viral particles stored in the VCC, via stimulation of the P2X7 purinergic receptor (Graziano et al., 2015). The rapid and drastic remodelling of the cell shape and actin cytoskeleton that eATP induces on HIV-infected MDMs may underlie its impact on viral release. This interpretation raises the question of how the structure of the VCC, their trafficking, and consequently viral particle release are impacted by or depend on the actin cytoskeleton.

The VCC are surrounded by a meshwork of filamentous actin tightly associated with its external membrane (Mlcochova et al., 2013; Pelchen-Matthews *et al.*, 2012), which is often decorated by an electron dense molecular coat that appears to anchor the actin cytoskeleton to the compartment (Pelchen-Matthews *et al.*, 2012). These coats have striking resemblances with focal adhesions, as they contain the ß2-integrin CD18 and its associated CD11b and CD11c α-integrins. Focal adhesion linker proteins, such as talin, vinculin and paxillin are also found at these molecular coats (Pelchen-Matthews et al., 2012). Silencing CD18 in HIV-1 infected macrophages, did not alter viral release over a 24-hour window. Whether a longer period of analysis impacts viral release was not assessed; nevertheless, CD18 silencing led to scattering of the VCC across the cell (Pelchen-Matthews *et al.*, 2012), suggesting that these coats are an integral part of the compartment.

In macrophages, many focal adhesion-mediated processes such as cell migration are regulated by Focal Adhesion Kinase (FAK)-related proline-rich tyrosine kinase 2 (PYK2) (Okigaki et al., 2003). PYK2 promotes the assembly of focal adhesions at the leading edge, and their disassembly at the trailing edge to ensure net forward movement (Zhu et al., 2018). The full breadth of PYK2 activators and substrates remain incompletely understood, but it is generally considered to act at the cross-roads between integrin and small GTPases, such as Rho (Schaller, 2010).

It remains unclear how the VCC structure and HIV particle release are impacted by these focal adhesion-like coats and PYK2 signalling and, in general, our understanding on how the actin cytoskeleton impacts HIV release from macrophages remains rudimentary. We report here that pharmacological stabilization of actin fibres led to scattering of the VCC and impeded viral particle release from infected MDM. This was paralleled by decreased association of the integrin CD18 and of the phosphorylated form of the kinase PYK2 with the compartment. Specific inhibition of PYK2 led to accumulation of viral particles in VCC that lost connection to the plasma membrane. Finally, subjecting infected MDM to frustrated phagocytosis on glass coverslip induced rapid trafficking of VCC to the surface and viral release. Such trafficking required the compartment to be associated with CD18 and the actin cytoskeleton. We propose that focal adhesion-like coats are regulated by PYK2 and anchor the actin cytoskeleton to the VCC to promote HIV release from macrophages.

## Results

### 1. Actin dynamics modulates viral particle release from HIV-1-infected macrophages

In MDMs infected with HIV-1-GAG-iGFP-ΔENV-VSVG for 4 days, phalloidin staining can be readily observed around the VCC, confirming previous observations (Mlcochova *et al.*, 2013) (Fig. 1A-B). F-Actin appears docked at or surrounding the compartment (see 3D inset in Fig. 1B). Ultrastructural analysis by electron microscopy (EM) revealed the presence of actin filaments anchored at and radiating from the VCC (Fig. S1A-B). Live imaging of MDMs transduced with Lifeact-mCherry and infected with HIV-1-GAG-iGFP-ΔENV-VSVG confirmed that the VCC and the actin cytoskeleton are in proximity to each other, and exhibit synchronized movements (Fig. 1C and Video S1).

**Figure 1.**
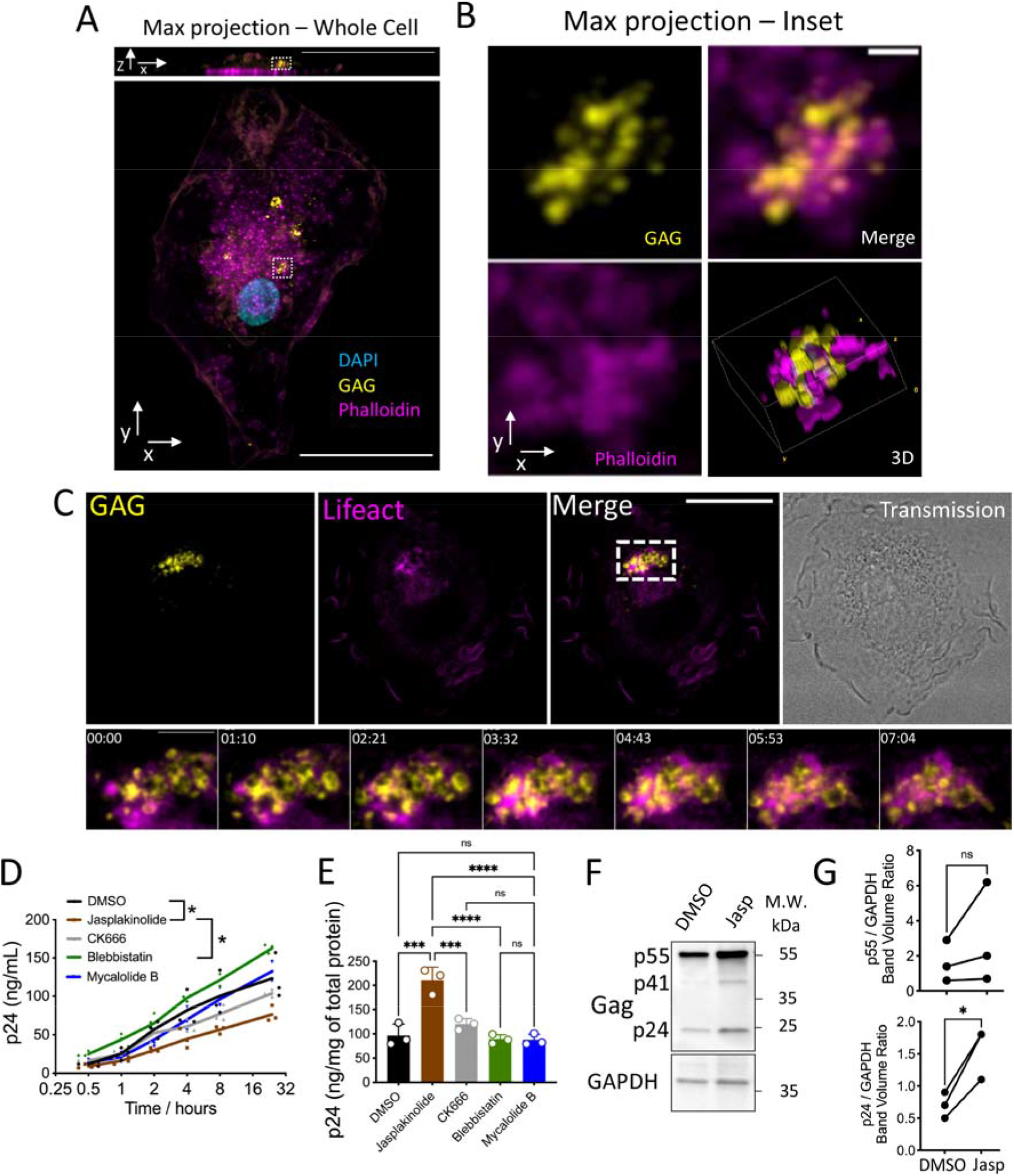
Actin is involved in VCC dynamics and viral release from macrophages. **A** – MDM infected with HIV-1 GAG-iGFP-ΔENV-VSVG for 7 days, stained for phalloidin. Scale bar = 30 μm. The bottom panel shows a maximum intensity z-projection. The top panel shows a slice on the xz plane, highlighting the selected region for panel 1B. **B** – Maximum intensity Z projection and 3D volume rendering of the region highlighted in 1A. Scale bar = 1 μm **C** – MDMs transduced with Lifeact-mCherry were infected with HIV-1-GAG-iGFP-dENV-VSVG (MOI=1.0). Time-lapse imaging was performed at day 6 after infection. The bottom panels represent the dashed region in the top panels. Scale bar= 20um (top panels) and 5um (bottom panels) **D** – MDMs were infected with HIV-1-ΔENV-VSVG at a MOI=1.0 for 4 days. Cells were subsequently treated with the indicated drugs for 24 hours. Supernatant aliquots were recovered at the indicated time-points, and the released p24 was quantified by a custom-made CBA assay. Jasplakinolide = 50 nM / CK666 = 10 μM / Blebbistatin = 5 μM / Mycalolide-B = 100 nM. Data from 3 independent donors. Two-way ANOVA. *= P<0.05 **E** – As in (D), cells were lysed at the end of the experimental procedure in NP-40 lysis buffer and intracellular p24 was quantified by CBA. Data from 3 independent donors. One-Way ANOVA – (** = P<0.01; *** P<0.001). **F** – As in (D), cells were lysed in RIPA buffer at the end of the experimental procedure and analysed by western blot for the expression of GAG. One representative donor is shown. **G** – Densitometry quantification of the western blot bands from (F) for 3 independent donors. Paired t-test, * = P<0.05, ns-non-significant

To explore how the actin cytoskeleton may impact on the structure of the VCC and viral release, we first employed a cell micropatterning technique to physically manipulate the actin cytoskeleton organization in HIV-1-infected MDMs (Fig. S2A-S2B). Indeed, adherent cells adapt to the micropattern shape by remodelling their actin cytoskeleton as to maximize adhesion (Thery, 2010). We observed that plating HIV-1-infected MDMs on distinct micropatterns impacted on the total volume of the VCC (Fig. S2A-S2B). This was particularly evident in crossbow-shaped macrophages, which presented stress fibres at the non-adherent edges, and bore significantly smaller VCCs (Fig. S2A-S2B). Nonetheless, these cells did not remain immobile over the patterned areas for prolonged periods of time, precluding any accurate analysis on viral particle release. We turned thus into pharmacological manipulation of the actin cytoskeleton of infected macrophages to ascertain how it impacts on the VCC size and structure as well as viral release.

We infected MDMs with HIV-1-ΔENV-VSVG for 4 days to allow formation of VCCs and treated them with a panel of pharmacological modulators of the actin cytoskeleton. Over the next 24 hours, we observed that the F-actin stabilizing drug, jasplakinolide, reduced the release of viral particles (Fig. 1D), as measured by our in-house CBA assay that specifically detects the HIV-1 p24 capsid protein (Fig. S3A). None of the other cytoskeleton-modifying drugs had significant effects on viral release over the 24 hours period (Fig. 1D). Jasplakinolide was also the sole drug to induce retention of p24 inside the macrophage during the 24 hours treatment period, suggesting that the viral particles are retained inside the cell, hence their decreased concentration in the supernatant (Fig. 1E-G). Interestingly, treatment of HIV-1-infected HeLa cells with the same cytoskeleton modulators failed to induce any significant changes in p24 release over vehicle-treated cells (Fig. S3B), suggesting that the effect of jasplakinolide treatment implicates the VCC, a macrophage-specific structure.

### 2. Stabilization of actin fibres impacts VCC structure and architecture

We next asked how jasplakinolide treatment results in retention of p24 in the macrophage, by examining its impact on the VCC by confocal microscopy. MDMs were infected with HIV-1-GAG-iGFP-ΔENV-VSVG for 4 days, treated or not with a non-toxic dose of jasplakinolide for 24 hours and analysed by confocal microscopy. As expected, the drug had a strong impact on the general macrophage morphology, as cells rounded up and lost their typical spreading over the substrate (Fig. 2A). While this led to a significant decrease in cell spreading (Fig. 2B), treatment with jasplakinolide increased the total volume of the VCC by more than 2-fold (Fig. 2C). The average intensity of the Gag-iGFP signal was similar between treated and not treated cells (Fig. 2D), although the compartment spread more evenly throughout the cell (Fig. 2E). These observations were confirmed by EM, as the typical VCC in vehicle-treated cells appear to fragment into smaller, but more numerous, compartments after treatment with jasplakinolide (Fig. 2F). Jasplakinolide-treated MDMs also exhibited massive accumulations of F-actin that occupied large sections of the cytoplasm (Fig. 2G, bottom left panel). Their appearance is concomitant with loss of the actin filaments that typically radiate from the VCC in control macrophages. Overall, these data suggest that drug-induced F-actin stabilization leads to loss of the organized actin filaments that are anchored at the VCC. Consequentially, compartments become dispersed throughout the cell and accumulate, resulting in impaired viral release.

**Figure 2.**
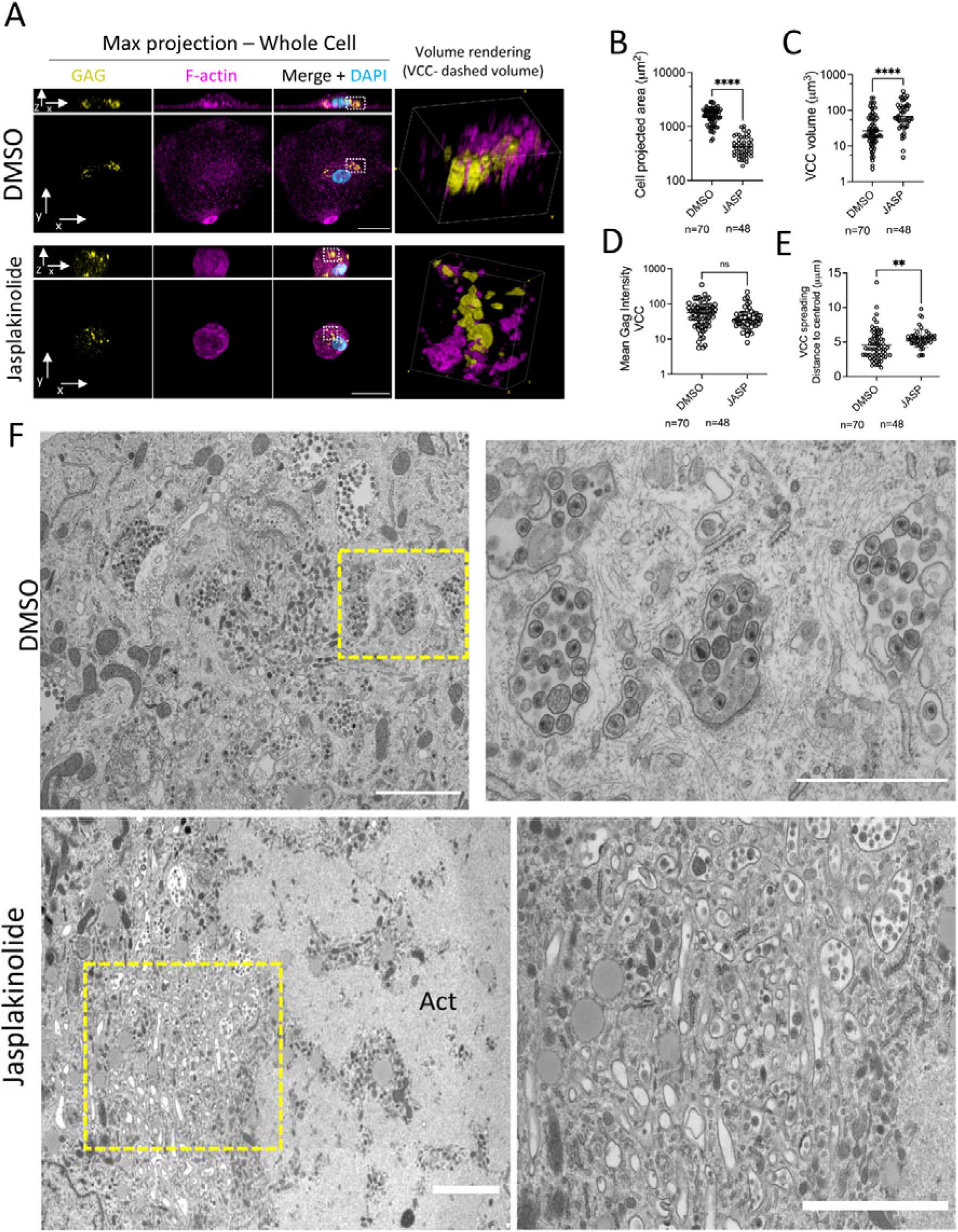
Stabilization of actin fibres scatters the VCC and increases the total volume of the compartment. **A** – MDMs were grown in coverslips, infected with HIV-1-ΔENV-GAG-iGFP-VSVG at a MOI=1.0 for 4 days and subsequently treated with DMSO or jasplakinolide (100 nM) for 24 hours. Cells were fixed, stained with anti-F-actin and imaged by confocal microscopy. **B-E** – Image analysis for cells processed as in (A) for the indicated parameters. See methods for details on the quantification strategies employed. Each circle represents one cell and the plots display cells from 3 independent donors. Unpaired t-test. (** = *P*<0.01; \*\*\*\**P*<0.0001). **F** – Cells were treated as in (A) and processed for electron microscopy. The right panels are enlargements of the areas enclosed in the dashed squares on the left panels. Scale bar = 2 μm (left panels), 1 μm (right panel).

### 3. Stabilization of actin fibres relocalizes the integrin CD18 away from the VCC-limiting membrane

Previous work reported that the β2-integrin CD18 is present in focal adhesion-like electron dense regions at the limiting membrane of the VCC and is critical for the integrity of the VCC (Pelchen-Matthews *et al.*, 2012). We confirmed the presence of electron dense regions that coat the VCC and that serve as anchoring sites for the actin cytoskeleton (Fig. 3A, Fig. 2G and Fig. S1A-B). These coats were also present in VCCs at, or close to, the cell surface (Fig. 3B), suggesting their implication on the trafficking of the compartment. Confocal immunofluorescence confirmed that CD18 surrounds the VCC (Fig. 3C-D). However, treatment with jasplakinolide lead to a significant decrease on the enrichment of CD18 at the vicinity of the VCC (Fig. 3C-D). Analysis by EM confirmed that the electron dense coats typical of control MDMs, were no longer observable in the smaller and more numerous VCCs of jasplakinolide-treated cells (Fig. 3E-F and Fig 2G). Our data indicates that pharmacological stabilization of actin leads VCCs to lose their association with focal adhesion-like coats and the integrin CD18, which possibly underlies the accumulation of viral particles inside the cell.

**Figure 3.**
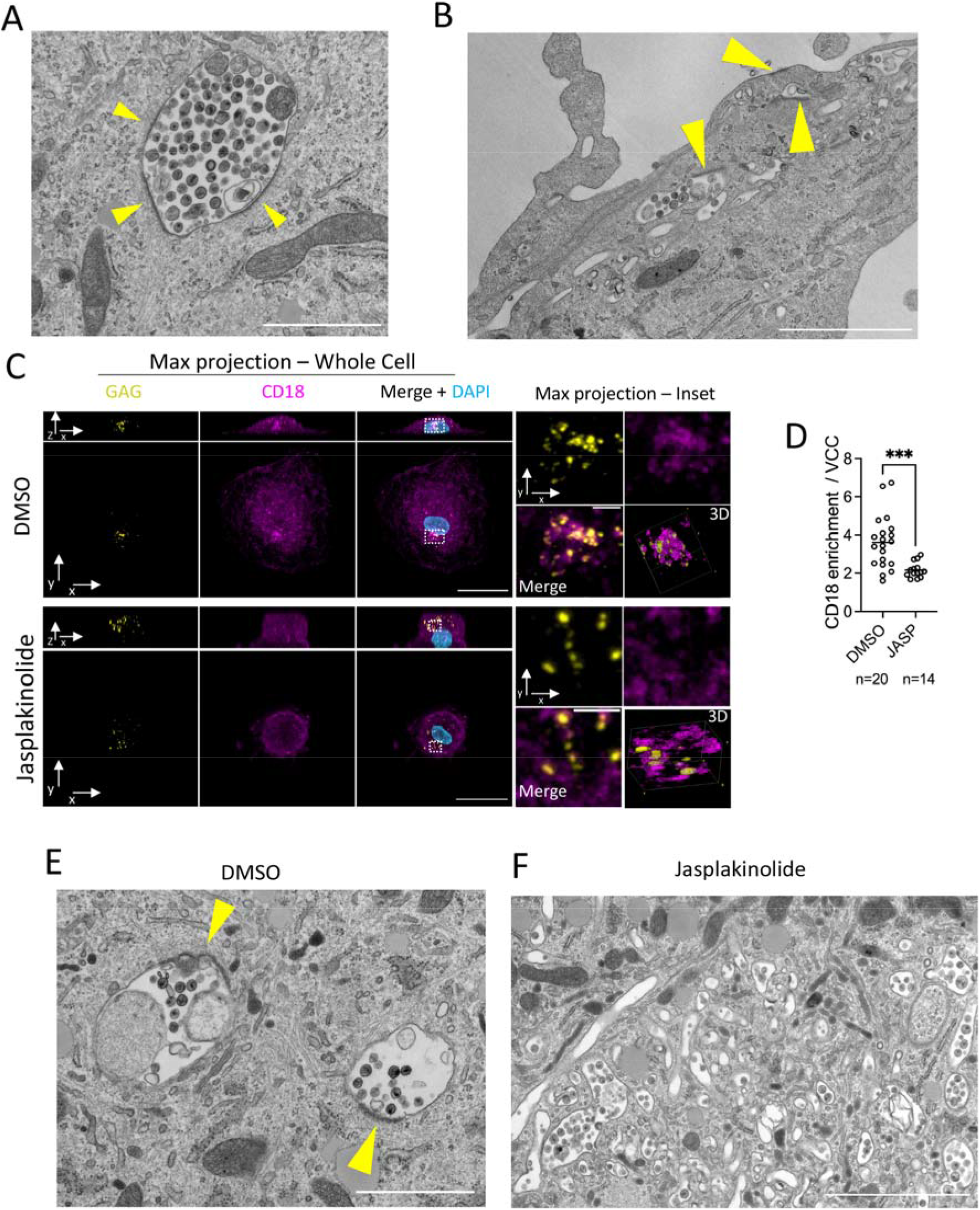
Stabilization of actin fibres relocalizes the integrin CD18 away from the VCC limiting membrane in HIV-1-infected macrophages. **A-B** – MDMs were infected HIV-1-ΔENV-VSVG (MOI=1.0) for 7 days. Cells were fixed and processed for electron microscopy. (A) depicts an example of an internal VCC with associated electron dense coats. (B) highlights the presence of compartments with coats close to the cell surface. Arrow heads point to the electron dense coats. Scaler bars= 2 μm. **C** – Confocal microscopy of MDMs infected with HIV-1-Gag-iGFP-ΔENV-VSVG (MOI=1.0) for 4 days and treated with jasplakinolide or DMSO for 24 hours. Whole cell projections are shown in the right, along the z-axis (bottom) or the y axis (top). On the left, the areas enclosed in the dashed squares are enlarged and shown as z-projections. Scale bar = 20 μm (left panels) or 2 μm (right panels). **D** – Quantification of CD18 enrichment at the VCC by image analysis. Each dot represents a single cell from 3 independent donors. See Methods for details on the analysis. **E – F** - Electron microscopy of MDMs infected with HIV-1-ΔENV-VSVG (MOI=1.0) for 4 days and treated with jasplakinolide or DMSO for 24 hours. Arrow heads point to electron dense regions associated with the VCC in DMSO treated cells (note their absence in cells treated with jasplakinolide). Scale bars= 2 μm.

### 4. The phosphorylated form of the focal adhesion kinase PYK2 localizes at the VCC

Because of the similarities between focal adhesions and the electron dense coats around the VCC, we evaluated whether PYK2, a non-receptor tyrosine focal adhesion kinase (FAK) that governs the focal adhesion response to integrin-mediated signalling, plays a role on the VCC structure and HIV-1 release from macrophages. A 4-day infection of MDMs with HIV-1-ΔENV-VSVG led to activation of PYK2, as evidenced by an increase in the levels of phosphorylated PYK2 at Tyr402 (Fig. 4A), in agreement with a previous report (Del Corno et al., 2001). Confocal microscopy revealed that P-PYK2 (Y402) localized mostly to the basal surface of the macrophage and was organized into large clusters that likely corresponded to podosomes (Fig. 4B). Yet, a small pool of P-PYK2 located at the VCC in DMSO-treated cells (Fig. 4B). While jasplakinolide treatment had no effect on the global levels of PYK2 Tyr 402-phosphorylation induced by HIV-1 infection (Fig. 4A), it led to loss of the enrichment of P-PYK2 at the VCC observed in vehicle-treated cells, as the P-PYK2 signal dispersed evenly throughout the cell (Fig. 4B-C). These data indicates that jasplakinolide treatment induces loss of association of the phosphorylated form of PYK2 from the VCC, accompanying the loss of CD18 and focal adhesion-like coats.

**Figure 4.**
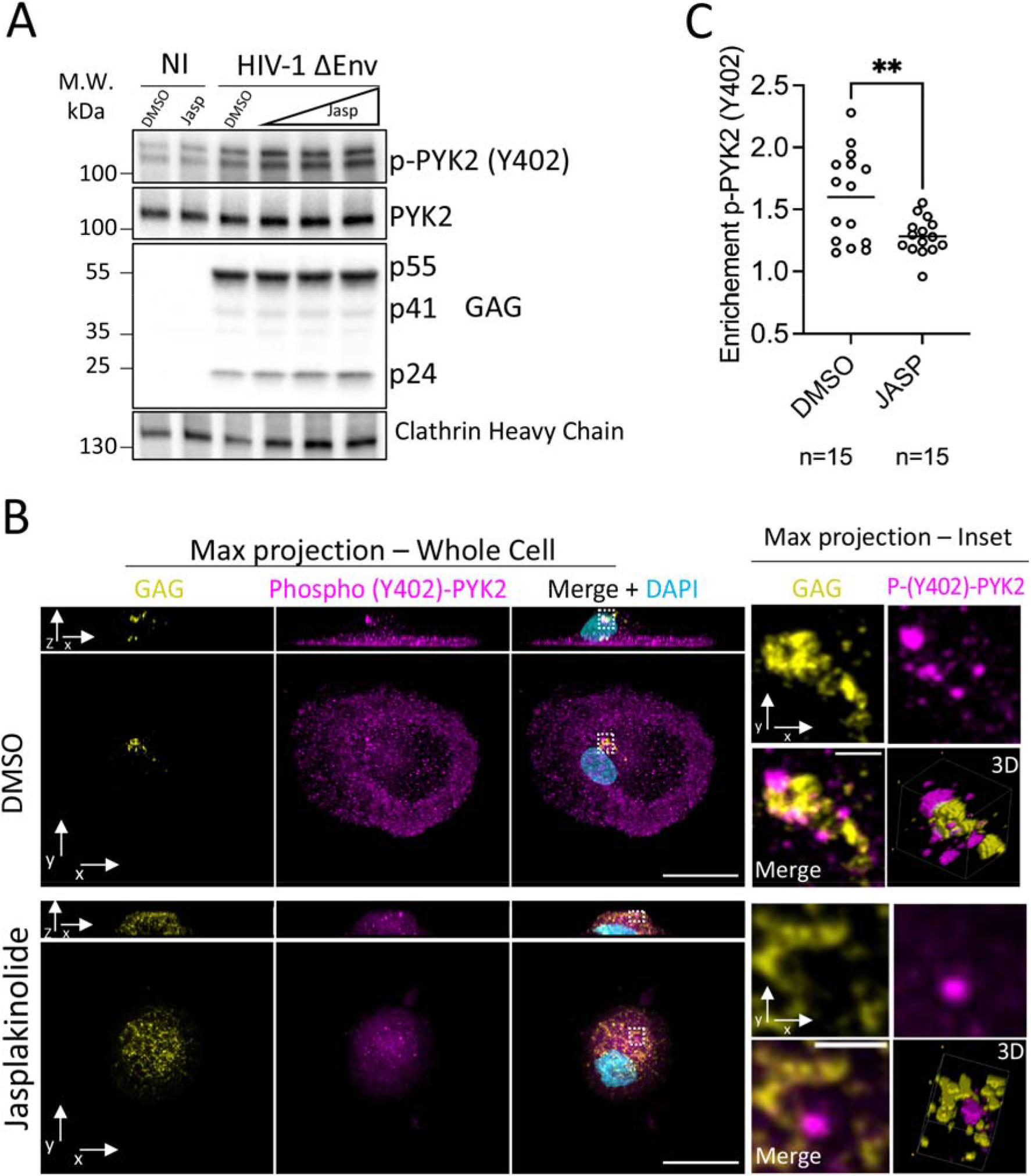
The phosphorylated form of integrin-associated kinase PYK2 partially colocalizes with the VCC. **A** – MDMs left uninfected or infected with HIV-1-Δ-ENV-VSVG (MOI=1.0) for 4 days were treated with jasplakinolide or vehicle (DMSO) for 24 hours. Cells were lysed and analysed by western blot for the indicated proteins. The higher concentration of Jasplakinolide employed was 200 nM and the lower concentrations correspond to consecutive two-fold dilutions. **B** – Confocal microscopy of MDMs infected with HIV-1-Gag-iGFP-ΔENV-VSVG (MOI=1.0) for 4 days and treated with jasplakinolide or DMSO for 24 hours. Whole cell projections are shown on the right, along the z-axis (bottom) or the y axis (top). On the left, the areas enclosed in the dashed squares are enlarged and shown as z-projections. Scale bar = 20 μm (left panels) or 2 μm (right panels). **C** – Quantification of P-(Y402)-PYK2 enrichment at the VCC by image analysis. Each dot represents a single cell from 2 independent donors. See Methods for details on the analysis.

### 5. Active PYK2 participates in viral particle release from the VCC

Overall, our results hint at a possible role for PYK2 as coordinating actin dynamics at the VCC, since (1) its active phosphorylated form localizes to the compartment, and (2) jasplakinolide treatment disrupts its localization at the VCC, concomitant with loss of the electron dense regions and CD18 expression at the compartment. To directly test this hypothesis, we treated HIV-1-ΔENV-infected MDMs with PF431396 (PF396), a dual FAK/PYK2 inhibitor that has higher affinity for PYK2 over FAK (IC50 values of 2 and 11 nM, respectively). PF396 prevented the phosphorylation of PYK2 induced by HIV-1 infection and decreased p24 release from infected macrophages, over a 96-hour period and, conversely, promoted its retention inside the cell (Fig. 5A-C). We next examined the VCCs of macrophages infected with HIV-1 GAG-iGFP-ΔENV-VSVG for 4 days and treated for additional 4 days with PF396 (Fig. 5D). While PF396 had no evident impact on the macrophage morphology, including cell volume (Fig 5E), it led to a significant increase in the compartment volume (from 42.26 μm^3^ in DMSO-treated cells to 103.13 μm^3^ in PF396-treated cells) (Fig. 5F), without an impact on the mean intensity of the Gag-iGFP signal (Fig. 5G). These VCC from PF396-treated MDMs often had a granular or reticulated appearance (Fig. 5D and S4A). While F-actin remained localized close to the compartment in PF396-treated cells, it appeared evenly distributed throughout the VCC, rather than being concentrated as patches in one side, as commonly observed in vehicle-treated cells (Fig 5D). Despite their larger volume, VCC from PF396 treated cells were more compact, as revealed by their decreased spreading scores (Fig 5H). Finally, while CD18 remained enriched in the vicinity of the VCC after PF396 treatment, its distribution was more even, as observed with F-actin (Fig. 5I and Fig. S4A).

**Figure 5.**
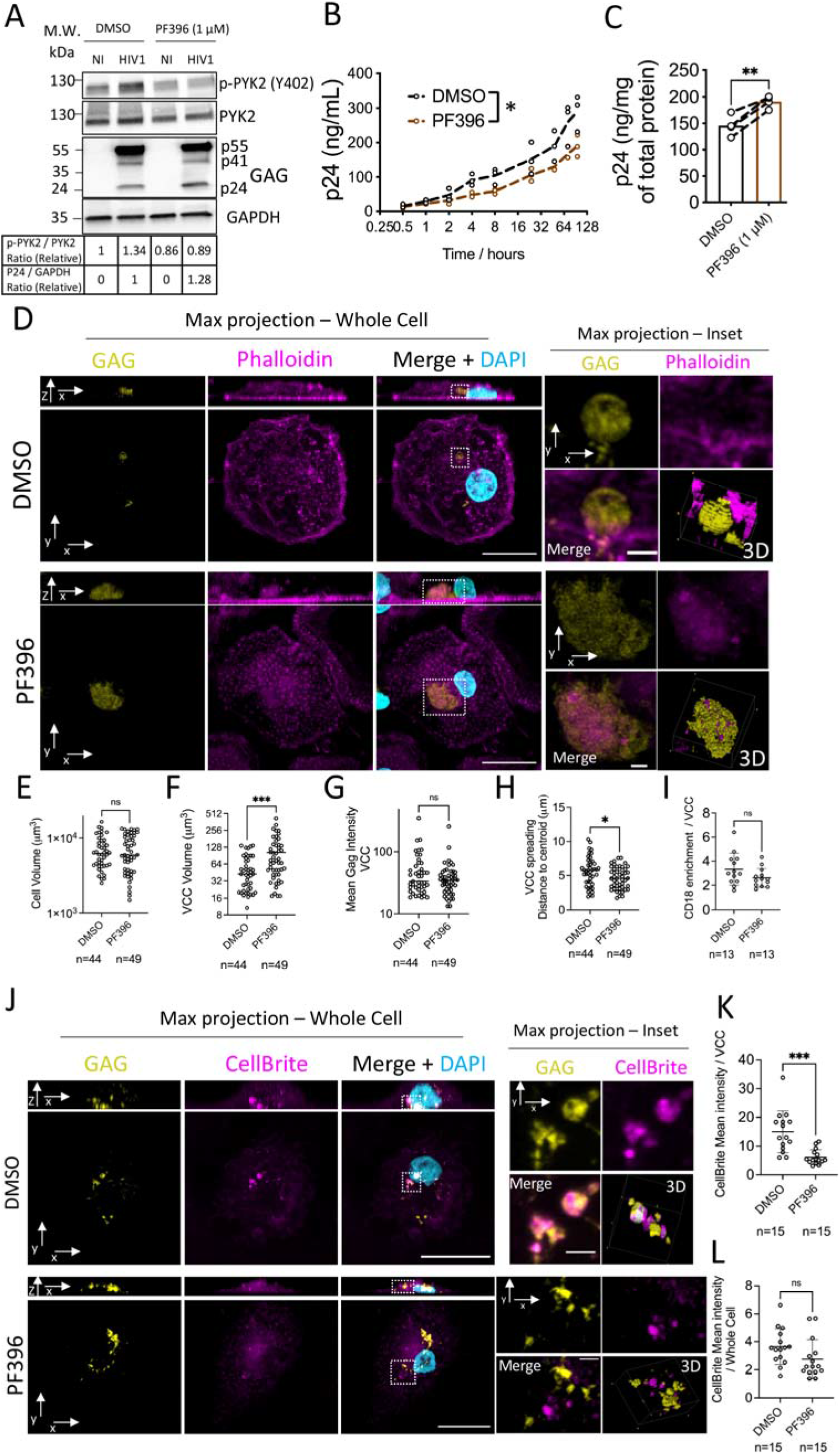
A PYK2 inhibitor promotes HIV-1 retention in VCC. **A** – MDMs left uninfected or infected with HIV-1-Δ-ENV-VSVG (MOI=1.0) for 4 days were treated with PF396 or vehicle (DMSO) for additional 96 hours. Cells were lysed and analysed by western blot with the indicated antibodies. The table below the blots display the intensity ratios indicated, as measured by densitometry. **B** – MDMs were infected with HIV-1-ΔENV-VSVG at a MOI=1.0 for 4 days. Cells were subsequently treated with PF396 (1μM). At the indicated time-points, supernatant aliquots were recovered, and the released p24 was quantified by CBA. Data from 3 independent donors. Two-way ANOVA. *= *P*<0.05 **C** – As in (B), cells were lysed at the end of the experimental procedure in NP-40 lysis buffer and intracellular p24 was quantified by CBA. Data from 3 independent donors. Paired t test – (** = *P*<0.01). **D** – Confocal microscopy of MDMs infected with HIV-1-Gag-iGFP-ΔENV-VSVG (MOI=1.0) for 4 days and treated with PF396 (1μM) or vehicle (DMSO) for an additional 96 hours. Whole cell projections are shown on the right, along the z-axis (bottom) or the y axis (top). On the left, the areas enclosed in the dashed squares are zoomed in and shown as z-projections. Scale bar = 20 μm (left panels) or 2 μm (right panels). **E-H** – Image analysis for cells processed as in (D) for the indicated parameters. See methods for details on the quantification strategies employed. Each circle represents one cell, and the plots show cells from 3 independent donors. Unpaired t-test. (* = *P*<0.05; *** *P*<0.001). **I** - CD18 enrichment at the VCC for cells processed as in Fig. S4. See methods for details on image analysis. Each circle represents one cell, and the plots display cells from 2 independent donors. Unpaired t-test. (ns= not significant). **J** – Confocal microscopy of MDMs infected with HIV-1-Gag-iGFP-ΔENV-VSVG (MOI=1.0) for 4 days and treated with PF396 (1μM) or vehicle (DMSO) for additional 96 hours. Cells were exposed to CellBrite before fixation. Right panel - whole cell projections along the z-axis (bottom) or the y axis (top). Left, panels – magnification of the areas enclosed in the dashed squares as z-projections. Scale bar = 20 μm (left panels) or 2 μm (right panels). **K-L** – Quantification of CellBrite intensity in the VCC (L) or elsewhere in the cell (M) for cells processed as in (K). See methods for details. Each circle represents one cell, and the plots show cells from 2 independent donors. Unpaired t-test. (Ns-non-significant; *** *P*<0.001).

Due to the increased compactness of VCC from PF396-treated macrophages, we hypothesized that the VCC continuity with the plasma membrane may be altered after exposure to the drug, which could explain the decreased viral particle release. We thus exposed PF396-treated or vehicle-treated macrophages to CellBrite®, a non-permeable lipophilic dye that can withstand fixation. Strikingly, we observed strong accumulation of the dye in VCC from DMSO-treated macrophages, while those from PF396-treated cells were generally negative for CellBrite staining (Fig. 5J-L and Fig. S5A). Finally, PF396 treatment did not render VCC accessible to an anti-CD44 antibody added from the extracellular media (Fig. S6A), although VCC from PF396-treated cells could still be stained with anti-CD44 after cell permeabilization (Fig. S6B). PYK2 is known for its critical role in regulating actin dynamics at focal adhesions (Schaller, 2010). Our data suggest that it is endowed with a similar role at the VCC, promoting its trafficking to the surface and regulating its connectivity to the plasma membrane.

### 6. Frustrated phagocytosis of HIV-1-infected MDM induces the rapid release of viral particles from VCC

While our previous experiments have examined the role of the actin cytoskeleton in regulating HIV release from resting macrophages, we aimed to observe its impact in cells undergoing a dynamic process. For that, we subjected HIV-1-infected MDMs to frustrated phagocytosis. In this experimental system, macrophages spread rapidly on the substrate, driven by actin polymerization that promotes a fast and concentric extension of pseudopods, until the plasma membrane tension increases to a point that no longer allows further spreading (Masters et al., 2013). If an HIV-1-infected macrophage is subjected to frustrated phagocytosis, we hypothesize that VCC act as membrane reservoirs that can be deployed rapidly to the surface to counteract the increase in plasma membrane tension, and that their anchoring to the cytoskeleton would be required for such rapid deployment. We thus seeded MDMs, infected for 3 days with HIV-1-GAG-iGFP-ΔENV-VSVG, over a human IgG-coated surface and performed live imaging. As exemplified in Fig. 6A, the macrophage spread rapidly on the surface with its projected area more than doubling in 20 minutes (Fig. 6A-B and Video S2). We further observed a steady decrease in the volume of the VCC starting at around 8 minutes (Fig 6A and 6B). The compartment appears to migrate towards the basal surface, as the cell spreads, where its intensity decreases, presumably due to virion release (Fig. 6A, xz planes). Indeed, by quantifying the amount of p24 in the culture supernatant, we observed a rapid release of the capsid protein from MDMs seeded over IgG-coated coverslips as compared with control coverslips or control coverslips + IgG added at the same time as the cells (“Soluble IgG”) (Fig. 6C), during the first two hours. Such rapid kinetics suggests that it is the preformed viral particles stored in VCCs that are released during frustrated phagocytosis and not a general increase in viral production due to macrophage activation. Supporting this idea, at later time-points there were no differences in the concentration of p24 released from macrophages seeded in the three different conditions (Fig. 6C). Thus, as macrophages spread during frustrated phagocytosis, VCC appear to be pulled to the basal surface of the cell, where they release their contents.

**Figure 6.**
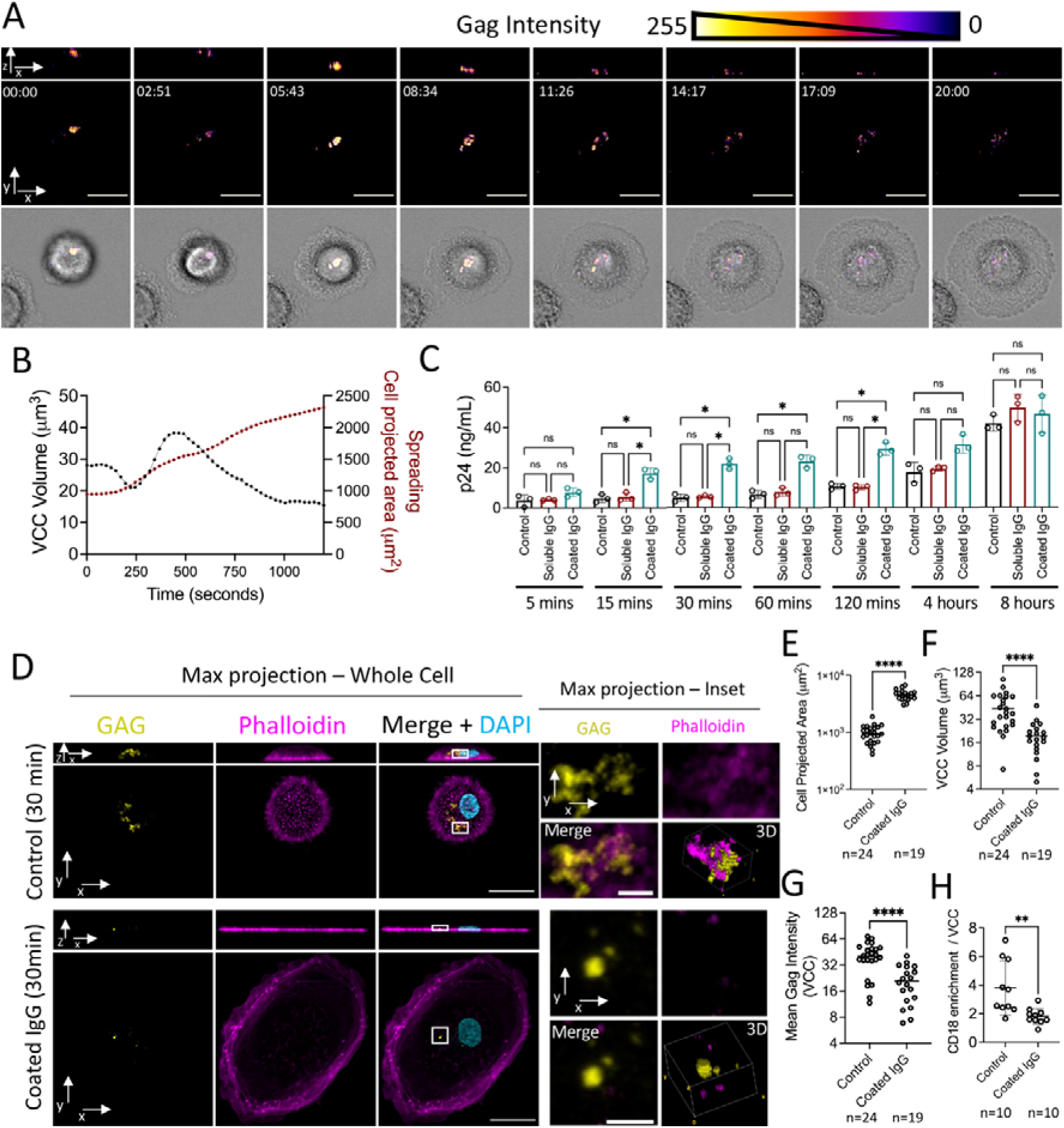
Rapid HIV release during macrophage frustrated phagocytosis is associated with trafficking of CD18+ VCC to the surface. **A** – Time-lapse imaging of MDMs infected with HIV-1-GAG-iGFP-ΔENV-VSVG for 3 days and seeded over human IgG-coats coverslips. Whole cell projections are shown for the top and middle panels, along the z-axis (middle) or the y axis (top). The bottom panel (transmission channel) is a single z-slice. Scale bar = 20 μm. **B** – Image analysis of the total VCC volume (left y-axis) and the cell projected area (right y-axis) over the course of the frustrated phagocytosis event depicted in (A). **C** – MDMs infected with HIV-1-ΔENV-VSVG for 3 days were seeded over human IgG-coated coverslips, or over non-coated coverslips, or in the presence of soluble IgG over non-coated coverslips. At the indicated time-points, aliquots of the supernatant were recovered and the released p24 was quantified by CBA. Data from 3 independent donors. For each time-point groups were compared by one-way ANOVA (* = *P*<0.05). **D** – Confocal microscopy of MDMs infected with HIV-1-Gag-iGFP-ΔENV-VSVG (MOI=1.0) for 3 days and seeded over control or human IgG-coated coverslips for 30 mins. Right panels - whole cell projections along the z-axis (bottom) or the y axis (top). Left, panels – magnification of the areas enclosed in the dashed squares as z-projections. Scale bar = 20 μm (left panels) or 2 μm (right panels). **E-H** – Image analysis for cells processed as in (D) for the indicated parameters. See methods for details on the quantification strategies employed. Each circle represents one cell, and the plots include cells from 2 independent donors. Unpaired t-test. (** = *P*<0.01; \*\*\*\**P*<0.0001).

We next tested whether the actin cytoskeleton participates in VCC trafficking during frustrated phagocytosis. We fixed HIV-1-infected MDMs subjected to frustrated phagocytosis or in control conditions and examined by confocal microscopy (Fig. 6D). In cells under frustrated phagocytosis, F-actin concentrated on the protruding edges and as expected, these cells spread over the glass surface (Fig. 6E). The VCC were significantly reduced in their volume in cells undergoing frustrated phagocytosis (Fig. 6F), as was the mean intensity of the Gag-iGFP signal (Fig. 6G). Interestingly, the remaining VCC in MDMs under frustrated phagocytosis were mostly devoid of associated F-actin (Fig. 6D). Furthermore, while VCC from control MDMs were associated with CD18, as expected, after frustrated phagocytosis the remaining VCCs lost this association (Fig. 6H and Fig. S7A). Finally, P(Y402)-PYK2 concentrated mostly in the protruding edges in cells under frustrated phagocytosis, while remaining VCC in general lacked P-PYK2 (Fig. S7B). This strongly suggests that the rapid deployment of the VCCs to the surface that occurs during frustrated phagocytosis, requires an associated cytoskeleton and focal adhesion-like coats, and that compartments lacking such associated complexes remain inside the macrophage after frustrated phagocytosis.

## Discussion

Overall, our data highlights the importance of the actin cytoskeleton and associated factors in regulating the release of HIV-1 from its VCC in macrophages, likely by providing a physical anchor that promotes its movement, connectivity with the plasma membrane and, consequentially, viral release. While previous studies addressed the presence of an actin cytoskeleton associated with the VCC, and its importance for viral release, the molecular details remained unknown. Here, we uncover an important role for the CD18-PYK2 axis as modulators of the VCC structure, its trafficking dynamics and viral release.

Cell shape is governed by the actin cytoskeleton, particularly cortical actin (Chalut and Paluch, 2016). To start exploring the importance of the actin cytoskeleton on HIV release from macrophages, we developed a micropatterning technique for HIV-1-infected macrophages. Our data indicates that the VCC’s structure and size are sensitive to cell shape. Particularly, macrophage adhesion to the crossbow pattern imposes strong physical constraints resulting in smaller VCC that are mostly devoid of surrounding F-actin. Accurate estimation of viral particle release was not achievable due to the strong adhesive properties of macrophages which tended to invade the surrounding anti-adhesive PEG-based substrate after a few hours of seeding over patterns. Nevertheless, our finding that PEG-Silane can, at least temporarily, efficiently micropattern primary human MDMs, may serve as a useful tool for future studies employing micropatterning to study human macrophage biology.

By exposing infected MDMs to different pharmacologic modulators of the cytoskeleton we found that the F-actin stabilizing drug jasplakinolide significantly impacted on viral release to the supernatant. A previous report demonstrated an increase in viral release from HIV-1-infected macrophages after treatment with the actin depolymerizing drug latrunculin, or the actin polymerization inhibitors, cytochalasin E or D (Mlcochova *et al.*, 2013). In our hands, the actin depolymerizing drug mycalolide B had no significant effect on viral release over a 24-hour period. Differences in the viral strains employed, experimental conditions or the distinct mechanisms through which mycalolide B and latrunculin depolymerize actin (Hayashi-Takanaka et al., 2019), may account for the divergent results.

The ARP2/3 complex inhibitor CK666 had no effect in viral release over 24 hours. Longer exposure to CK666 led to toxic effects, so we cannot exclude that in the long-term actin polymerization may be required for viral release. Nevertheless, at least in the short term, actin polymerization may not be necessary for HIV release from macrophages. Similarly, the myosin II inhibitor blebbistatin had no significant effect on viral release over a 24-hour period, suggesting that actomyosin contraction is not a major driver of viral release, at the steady state.

The use of pharmacological manipulation of the actin cytoskeleton is not without its shortcomings, due to its critical importance in multiple other biological processes. During HIV infection actin impacts virtually on every step of the viral cycle (Ospina Stella and Turville, 2018). To minimize “off-target” effects, we allowed the establishment of the infection and formation of VCCs for 4 days before addition of the cytoskeleton modulators, as to exclude any effect on the early steps of the viral cycle. Nevertheless, we can’t dismiss certain unwanted effects may indirectly impact on HIV release.

Jasplakinolide treatment had no effect on p24 release from infected HeLa cells. Previous work showed that while latrunculin augmented viral release from macrophages, it had no effect on infected HEK 293 cells (Mlcochova *et al.*, 2013). Given that in these cell lines HIV-1 viral particles directly bud to the extracellular media, their release may not be impacted by modulating actin dynamics. Nevertheless, studies addressing the impact of viral release from non-macrophage cells often yielded inconsistent results (Ospina Stella and Turville, 2018). Some reports suggest that actin may be important for the assembly and budding processes (Thomas et al., 2015; Wen et al., 2014), but this is at odds with a study that tracked individual budding sites in real-time in HeLa cells, demonstrating that assembly and budding proceed unimpaired even after total depolymerization of actin (Rahman et al., 2014). Thus, it will be important to standardize experimental approaches to study the effects of the actin cytoskeleton on HIV assembly and budding. In macrophages, the field will face the added challenge of discriminating between the impact of actin modulation on viral assembly and budding, and its influence on VCC trafficking, as both may impact on viral release.

In HIV-infected macrophages exposed to jasplakinolide, VCC scattered throughout the cell and were smaller, but more numerous, such that their total volume increased. They were devoid of associated actin, CD18 and the focal adhesion-like coats typically found in non-treated infected cells. In non-treated and infected macrophages, we observed that actin bundles frequently radiated from these VCC coats, suggesting that they are the site of anchoring of the cytoskeleton to the compartment. Jasplakinolide induces the loss of these structures, concomitant with the fragmentation of the compartment, suggesting an important role for the coats and their associated cytoskeleton in maintaining the integrity of the VCC.

Our frustrated phagocytosis assays indicate an important role for the actin cytoskeleton and VCC-associated CD18 in the trafficking of the compartment to the surface. We show that the drastic remodelling of the macrophage morphology that occurs in frustrated phagocytosis leads to rapid viral release, associated with loss of VCCs. Importantly, those VCC remaining in the cell after frustrated phagocytosis were small and mostly devoid of actin and CD18, suggesting that these are required for rapid VCC trafficking to the surface and viral release. VCC appear to be pulled rapidly to the basal surface of the cell during frustrated phagocytosis. Is the pulling force transmitted through the actin bundles connected to the compartment, as our observations suggest? If yes, where does such force originate from, and what is the role of the electron dense coats that surround the VCC? One may speculate that the coats act as traction points, upon which pulling forces may act, in a manner akin to focal adhesions during mesenchymal cell migration (Bear and Haugh, 2014). In this scenario, one would expect a role for actomyosin contraction at the origin of the pulling forces acting on the VCC. While we failed to observe any effect of myosin II inhibition in viral release from macrophages at the steady state, its impact may be more prominent during an event such as frustrated phagocytosis, where VCCs rapidly traffic through the cell. However, given the known role for myosin II in the formation of phagocytic cups, we could not test the impact of its inhibition in viral release during frustrated phagocytosis (Olazabal et al., 2002). Future studies, employing alternative approaches able to specifically isolate the role of myosin II in VCC trafficking could help answer this question.

We found that the phosphorylated form of the kinase PYK2 localizes at the VCC, and that jasplakinolide treatment redistributes it throughout the cell. The key role this kinase plays in regulating the dynamics of focal adhesions in macrophages, prompted us to explore its importance in HIV release. Inhibiting PYK2 led to viral particle accumulation in the macrophage, associated with larger VCC that often had a granular appearance. While F-actin and CD18 associated with the VCC via patches or focal points in control cells, treatment with the PYK2 inhibitor led to a more even distribution of these factors on the compartment. This suggests that PYK2 inhibition leads to altered organization of the focal adhesion-like coats that attach the actin cytoskeleton to the VCC, which may possibly underlie its impact on viral release. PYK2 inhibition also led VCC to lose plasma membrane connectivity, concomitant with the increased compaction of the compartment and perinuclear location. In vehicle-treated and infected cells, certain VCC were connected to the plasma membrane, while others not, in agreement with previous observations (Deneka et al., 2007; Gaudin et al., 2013). In bone marrow macrophages from humanized mice infected with HIV-1, completely enclosed compartments were often perceived, and these occasionally fused with plasma membrane-connected compartments (Ladinsky *et al.*, 2019). Integrating these observations with our new data, VCC appear to alternate between enclosed and plasma membrane-connected states, in a manner regulated by PYK2 activity. These enclosed compartments maintain the expression of VCC classical markers such as CD18 and CD44, despite remaining inaccessible to antibodies added prior to cell fixation and permeabilization.

PYK2 phosphorylation has been previously reported during HIV-1 infection and occurs downstream of co-receptor engagement by the envelope glycoprotein gp120 (Del Corno *et al.*, 2001), but not when Δ*Env*, pseudotyped viruses are used (Davis et al., 1997). Thus, our data points to a distinct, lasting, activation of PYK2 during HIV-1 infection mediated by an undescribed mechanism. In macrophages, HIV can reprogram cell migration, via its accessory protein Nef, towards a mesenchymal mode (Verollet et al., 2015), which depends on adhesive structures such as podosomes (Van Goethem et al., 2010) and is associated with PYK2 activation (Duong and Rodan, 2000). It is thus possible that a later activation of PYK2 by HIV Nef may contribute to enhancing viral release from macrophages.

A limitation of our study that could be explored in future research was the use of *Env*-deficient viruses pseudotyped with VSV-G to achieve single-cycle infections. This strategy avoids propagation of the infection, which could indirectly impact the quantification of viral release. However, in our cultures, direct viral transmission from one cell to another does not occur due to the absence of the viral envelope. VCC are often observed in proximity to the viral synapse that forms between an infected and a target cell (Duncan *et al.*, 2014; Gousset et al., 2008), where they potentially participate in direct viral transmission. Interestingly, direct transmission of HIV-1 from macrophages to CD4^+^ T cells is inhibited by jasplakinolide (Duncan *et al.*, 2014), suggesting that the same actin-dependent mechanisms that regulate VCC trafficking for viral release, also impact its redistribution towards viral synapses. It would be valuable to explore whether the factors described in this work as regulating viral release and VCC trafficking, such as PYK2 or the VCC coats, also play a role in cell-to-cell HIV transmission.

To conclude, whether at the steady-state or upon a stimulus that requires sudden and drastic changes in macrophage morphology, VCC appear to be dynamic structures that traffic through the cell in a manner that depends on its associated actin and microtubule cytoskeleton (Gaudin et al., 2012). In tissues, macrophages are constantly subjected to mechanical constraints as they navigate through crowded tissues, in a manner dependent on cytoskeletal dynamics. Our work suggests that HIV may exploit the macrophage biology to promote its own release and propagation.

## Material and Methods

### Cells

Plasmapheresis residues from healthy adult donors were obtained from the Établissement Français du Sang (Paris, France). All donors signed informed consent allowing the use of their blood for research purposes. Peripheral blood mononuclear cells (PBMC) were separated using Ficoll-Paque (GE Healthcare), and monocytes were isolated by positive selection using CD14+ magnetic beads (Miltenyi Biotec) and differentiated into macrophages for 7 days in RPMI (Gibco, Life Technologies) supplemented with 5% fetal calf serum (FCS; Gibco), 5% human serum AB (Sigma), penicillin-streptomycin (Gibco), and 25 ng/ml macrophage colony-stimulating factor (M-CSF; Miltenyi Biotec). HeLa and HEK 293 FT cells were maintained by bi-weekly subpassage in DMEM (Gibco, Thermofisher Scientific), supplemented with FCS and antibiotics.

### Virus production and monocyte transduction with LifeAct-mCherry

Viral particles were produced by transfection of 293FT cells in 6-well plates with 3 μg of total DNA and 8 mL TransIT-293 Transfection Reagent (Mirus Bio) per well. Plasmid mixes to produce HIV-1 viral particles consisted of 0.4 μg CMV-VSVG (pMD2.G; 12259; Addgene) and 2.6 μg of HIV proviral plasmid. To quantify viral release we preferentially used viral particles produced from pNL-ΔENV-VSVG which was kindly provided by O. Schwartz (Institut Pasteur, Paris, France). For examination of the VCC by microscopy we employed the pNL-GAG-iGFP-ΔENV-VSVG, a viral construct in which EGFP is placed between viral protease cleaving sites and between the matrix and capsid domains of Gag (Hubner et al., 2007). Media was renewed 16 hours later, and viruses were harvested at around 48 hours after transfection, filtered through a 0.45 μm pore and stored at −80°C until use. Thawed viral stocks were titrated by infecting the GHOST X4R5 reporter cell line.

A similar protocol was followed to produce lentiviral particles for monocyte transduction with Lifeact m-cherry. Here, transfection mixes consisted of 0.4 μg CMV-VSVG, 1.0 μg of the packaging plasmid PSPAX2 (12260; Addgene) and 1.6 μg of LifeAct-mCherry lentivector (pCDH1-CMV-LifeAct-mCherry). In parallel, VPX-containing SIV particles were produced by transfecting 293FT cells with 0.4 μg CMV-VSVG and 2.6 μg of pSIV3+ plasmid (Mangeot et al., 2000). Monocytes were transduced with equal volumes of freshly harvested LifeAct-mCherry and SIV3 lentiviral particles in the presence of 8 μg/mL of protamine (Sigma). At 48 hours, differentiating MDMs were exposed to 2 μg/mL of puromycin to select for successfully transduced cells, and were harvested for experiments at day 7 of differentiation.

### *In vitro* infections and cell treatments

MDMs were infected with HIV-1 NL-GAG-iGFP-ΔENV-VSVG or NL-ΔENV-VSVG at a MOI of 1.0. After 16 hours, culture media was renewed, and the infection allowed to progress for further 72 hours. At day 4 after infection, cells were exposed to the drugs or vehicle (DMSO) at concentrations and for the time periods indicated in figure legends. Jasplakinolide (Sigma); CK666 (Sigma); Blebbistatin (Sigma); Mycalolide B (Enzo Life Sciences); PF431396 (Tocris).

HeLa cells were infected with NL-ΔENV-VSVG at a MOI=1.0. After 16 hours, the culture media was renewed, and the same drugs added at similar concentrations.

To assess the membrane connectivity of the VCC, we employed the CellBrite Fix membrane stain. Briefly, infected MDMs were rinsed in PBS and incubated for 15 min at 37°C with a 1/1000 dilution of CellBrite Fix-640 (Biotium). Cells were rinsed twice with PBS and fixed in PFA 4% for confocal microscopy.

To assess the accessibility of VCC to anti-CD44 antibodies, infected MDMs were rinsed in PBS and stained with anti-CD44 antibodies at RT for 15 mins, before rinsing twice with PBS and fixation with PFA 4%. Cells were subsequently processed for confocal microscopy as described below.

### Macrophage Micropatterning

We employed a deep UV light-based method to imprint micropatterns onto glass coverslips. 12-mm coverslips were plasma treated and incubated with a solution of 0.1 mg/mL of poly(acrylamide)- μg - (PEG, 1,6-hexanediamine,3-aminopropyldimethylethoxysilane, (PAcrAm-g-(PEG, NH2, Si), Surface Solutions, Zurich) (Serrano et al., 2016) in 10 mM HEPES pH=7.4. A quartz mask containing the transparent motives for micropatterning was thoroughly washed with 70% ethanol, and coverslips were attached to its quartz surface using a small volume of water. The mask was then subjected to deep UV illumination in an UV ozone oven (UVO cleaner, model 342-220, Jelight) for 10 min into its silver face to induce degradation of PEG chains in the coverslips accessible through the transparent motives. Coverslips were finally detached from the mask and incubated for 1 hour at RT with a 5 μg/mL solution of fibronectin (Sigma) in NaHCO3 100 mM. MDMs, previously infected with HIV-1 NL-GAG-iGFP-ΔENV-VSVG (MOI=1.0) for 3 days, were seeded over patterned coverslips (20000 cells per coverslip) in pre-warmed media and allowed to attach for 1 hour at 37°C. Non-adherent cells were washed by media changes and the adherent cells were fixed in PFA 4%.

### Frustrated Phagocytosis

Twelve mm glass coverslips were extensively washed with 70% ethanol, dried and incubated with 10 μg/mL of IgG from human serum (Sigma) in PBS for 1 hour at RT. For quantification of p24 release, 24-well culture plates were directly coated with human IgG. MDM were infected with HIV-1 NL-GAG-iGFP-ΔENV-VSVG for microscopy, or with HIV-1 NL-ΔENV-VSVG for p24 quantification at a MOI=1.0. At day 3 after infection, cells were detached, washed and seeded over IgG-coated substrates (5000 cells for microscopy, 20000 for p24 quantification). Cells were subsequently fixed in PFA at 30 mins for fixed cell confocal microscopy, or immediately imaged for live cell imaging. For p24 release quantification, aliquots of the supernatant were collected at the indicated time-points, filtered (0.45μm) to remove floating cells, and stored until quantification by the p24 CBA.

### Quantification of p24 in culture supernatant or cell lysates by cytometric bead array (CBA)

Culture supernatant or cell pellets were lysed in a buffer containing 0.1% BSA, 1% NP-40 and 0.02% Tween-20. The amount of p24 in samples was subsequently quantified by a custom-made CBA assay that detects the HIV-1 p24 protein. Briefly, B6 functional beads (BD Biosciences) were conjugated with an anti-GAG antibody (H183-H12-5C hybridoma (mouse IgG1), NIH) and serve as capturing beads by incubation with samples for 1 hour at room temperature with agitation. An anti-p24 detection antibody (KC57-FITC, Beckmann Coulter) is subsequently added and incubated for 2 hours at RT. After two washes in a buffer containing 0.1% NP-40, beads are flowed in a FACS Verse (BD Biosciences) and the p24 concentration in culture estimated using a standard curve with serial dilutions of recombinant p24. For cell lysates an aliquot of the sample was used to quantify total protein by the BCA method (Pierce Micro BCA kit, Thermo Fisher Scientific) and the data is presented as μg of p24 / mg of total cell protein.

### Western Blot

Cells were lysed in RIPA buffer and total protein levels quantified with the BCA method. 30 μg of sample was reduced with Laemmli buffer, boiled and loaded in pre-cast polyacrylamide gels (BioRad). After transfer onto PDVF membranes (BioRad), blocking buffer (5% non-fat dry milk (m/v) in PBS-Tween-20 0.1% (v/v)) was added for 1 hour at RT. Primary antibodies were added overnight at 4°C with gentle agitation in antibody dilution buffer (5% BSA in PBST), washed and incubated with species-matched secondary HRP-conjugated secondary antibodies for 1 hour at RT. Membranes were revealed in a BioRad Chemidoc after addition of HRP substrate (Clarity, Biorad). Densitometry quantifications were performed using the ImageLab software (BioRad).

### Confocal microscopy and Immunostaining

Cells were fixed in 4% paraformaldehyde (PFA), washed in PBS and then permeabilized with permeabilization buffer (BSA 1% (m/v), Triton X-100 0.3% (v/v) in PBS) for 1 hour at room temperature. The primary antibodies used were anti-F-actin, anti-CD18, anti-pPYK2 and anti-CD44, as listed in supplementary table 1. Primary antibodies were diluted in permeabilization buffer, and the staining was performed overnight at 4°C in the dark. Subsequently, cells were washed 3 times and secondary staining was done in permeabilization buffer supplemented with 5% Donkey serum (Sigma) for 1 hour. For secondary antibodies, donkey anti-rabbit A647 or donkey anti-mouse Alexa 647 (Invitrogen) were used. Phalloidin-AF647 (Invitrogen) staining was also performed at this stage. Cells were washed 3 times then rinsed in PBS and mounted in Fluoromont-GTM medium with DAPI (Invitrogen).

Cell imaging was performed on an inverted confocal microscope (Leica DMi8, SP8 scanning head unit) equipped with a 63x oil immersion objective (NA=1.4) and four laser diodes (405, 488, 546, and 633 nm). An optimized z-step of 0.31 μm was employed for all acquisitions.

### Image analysis and quantification

Image J was used for all image analyses and quantifications, using custom-made macro scripts to automate the analysis. The Gag-iGFP signal was used to define a binary mask on the VCC, at each z-slice. The total volume of the VCC was then obtained by summation of the binary mask value, at each x,y pixel, across all the z-stack, correcting for the known dimensions of the voxel. A similar strategy was employed to estimate the total cell volume. Depending on the staining, the binary masks to define the cell area at each z-slice were derived from the phalloidin, F-actin or CD18 staining. Mean Gag intensity corresponds to the average Gag-iGFP signal at the region defined as the VCC by the binary mask.

To calculate VCC dispersion (spreading) across the cell, we used maximum intensity projections of the VCC’s binary mask and determined its centroid. Spreading was then determined by calculating the average distance of each VCC’s pixel in the projected mask to the centroid. A similar strategy was used to calculate the average distance of the VCC to the nuclear centroid. In this case, DAPI was used to define the nuclear binary mask, which after maximum intensity projection, was used to determine its centroid.

To determine the enrichment of CD18, p-PYK2 or CellBrite at the VCC, we first defined the VCC and cell binary masks and obtained the average intensity for the CD18, p-PYK2 or CellBrite staining in both the VCC and the whole cell. The enrichment score was then obtained as the ratio between the average intensity at the VCC and the average intensity in the whole cell.

All image montages were assembled using ImageJ. Y-projections were obtained using the Reslice tool, followed by maximum intensity projections. The 3D reconstructions which were achieved by applying the Volume Viewer plugin to the binary masks of the channels imaged.

### Live cell imaging

For live imaging experiments, cells were imaged on an inverted confocal microscope (Leica DMi8, SP8 scanning head unit) equipped with a 40X oil immersion objective and a live cell imaging chamber to maintain a constant temperature of 37°C and CO2 levels of 5%.

To assess the dynamics of the actin cytoskeleton surrounding the VCC, MDMs transduced with lifeact-mCherry were infected with HIV-1 GAG-iGFP ΔENV-VSVG at a MOI=1.0 for 6 days in glass bottom 35-mm dishes (Ibidi). Z-stacks were acquired in real-time at a frame rate of 0.42/s.

To image infected MDMs during frustrated phagocytosis, cells were placed in the imaging chamber immediately after their addition to IgG-coated 35-mm glass bottom disks. Z-stacks were acquired at a frame rate of 0.041/s. Acquired images were processed in ImageJ to assemble videos and montages.

### Electron microscopy

MDMs infected with HIV-1-NL-ΔENV-VSVG, were fixed in 2% glutaraldehyde in 0.1 M cacodylate buffer, pH 7.4 for 1 hour and subsequently fixed for 1 hour in 2% buffered osmium tetroxide, dehydrated in a graded series of ethanol solution, and then embedded in epoxy resin. Images were acquired with a digital camera Quemesa (SIS) mounted on a Tecnai Spirit transmission electron microscope (FEI) operated at 80 kV.

### Statistical analysis

Data were analysed using Prism software v9 (GraphPad). If not specified in figure legends, graphics show individual donors as dots with bars or lines at the means. Distributions were assumed to be non-parametric, and the specific test employed is indicated at the figure legend. ns, P value of >0.05; *, P value of ≤0.05; **, P value of ≤0.01; ***, P value of ≤0.001; ****, P value of ≤0.0001.

## Acknowledgements

We thank Dr Claire Hivroz and Ana-Maria Lennon-Dumenil at Institut Curie for fruitful discussions and critical reading of the manuscript. We also thank François-Xavier Gobert and Nicolas Carpi for discussions or technical help. This work was supported by grants from Agence Nationale de Recherche contre le SIDA et les hépatites virales (ANRS), Ensemble contre le SIDA (Sidaction), Laboratoire d’Excellence (Labex) DCBIOL (ANR-10-IDEX-0001-02 PSL and ANR-11-LABX-0043) to P.B. V.R. was supported by fellowships from ANRS and Sidaction.

## Author Contributions

VR, PB and MM designed the research. VR, ST, MSR, EG and AH performed the research. VR, ST, MM and MSR analysed the data. VR and PB supervised research. VR, ST and PB wrote the paper.

