## Supplementary figures and tables for "Release of HIV-1 particles from the viral compartment in macrophages requires an associated cytoskeleton and is driven by mechanical constraints"

**Supplementary materials.**

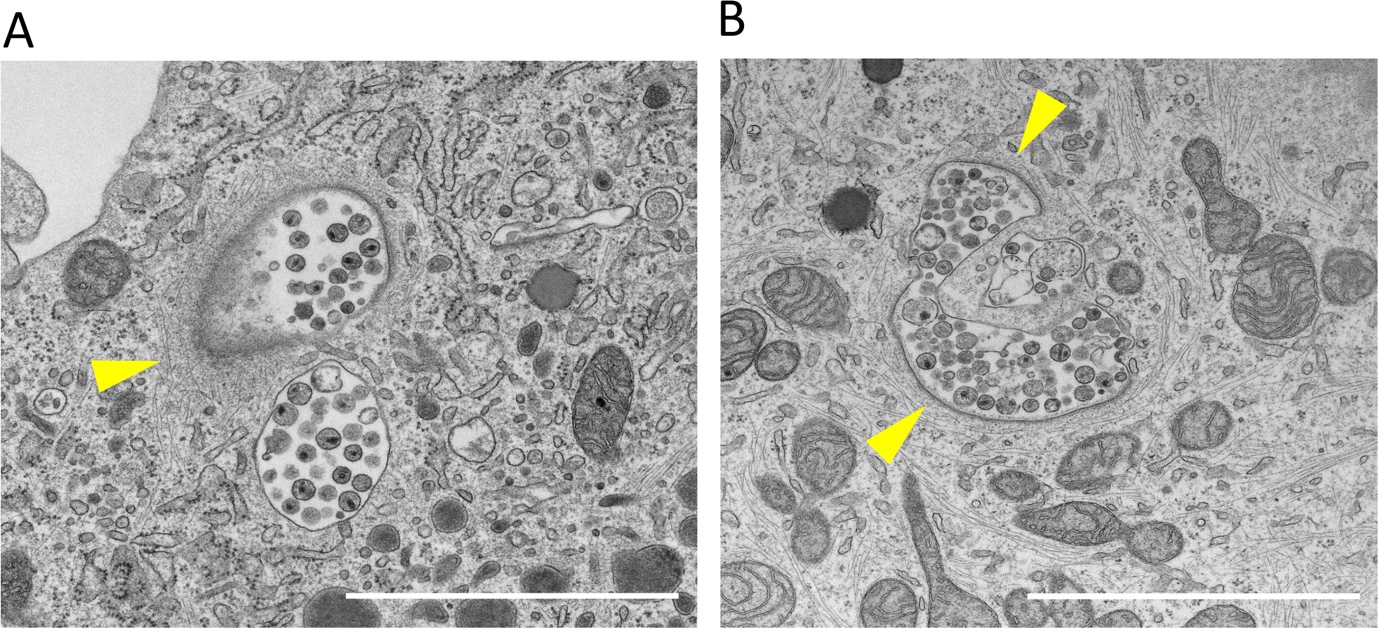

**Figure S1**. Actin is tightly associated with VCC at the ultrastructural level

**A-B** – MDMs were infected with HIV-1-ΔENV-VSVG (MOI=1.0) for 7 days. Cells were fixed and processed for electron microscopy imaging. Arrowheads depict the VCC-associated actin cytoskeleton. Scale bar = 2 µm.

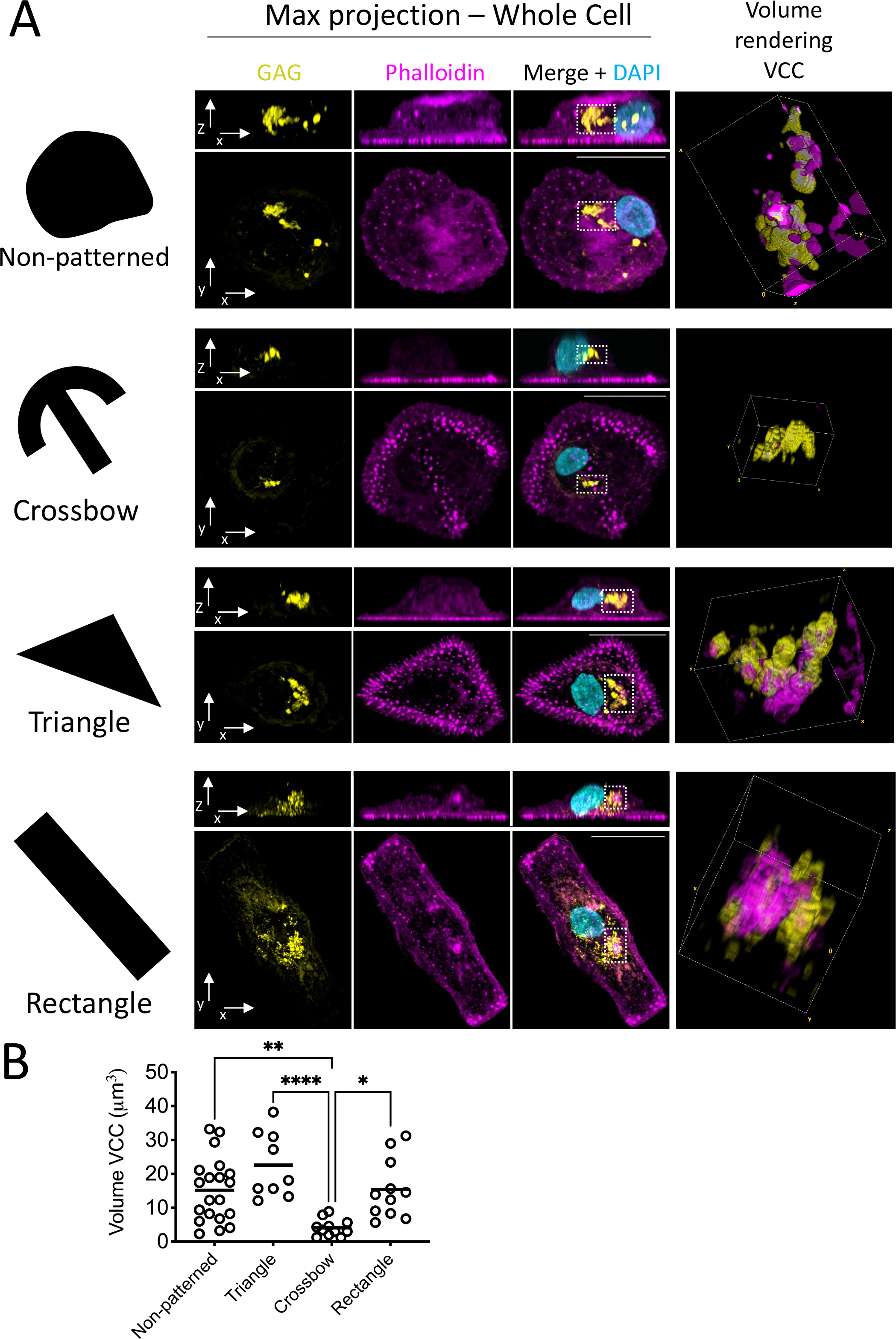

**Figure S2**: Physical constraints induced by micropatterning of HIV-1-infected macrophages impact VCC volume

**A** – MDMs were infected by HIV-1-GAG-iGFP-ΔENV-VSVG (MOI=1.0) for 3 days and seeded over coverslips bearing micropatterns with the indicated shapes. Cells were fixed 2 hours after seeding and stained for confocal microscopy. For each pattern type, top panels represent maximum projections in the y-axis, while the bottom panels are maximum projections across the z-axis. The 3D rendering at the right is an enlargement of the volume delimited by the dashed areas. Scale bar = 20um

**B** – The VCC volumes for each individual cell were estimated using image analysis of the Gag signal intensity. Each dot represents the total VCC volume within a single cell from 2 independent donors. Groups were compared using one-way ANOVA, followed by Tukey’s multiple comparisons test. P<0.05(*)

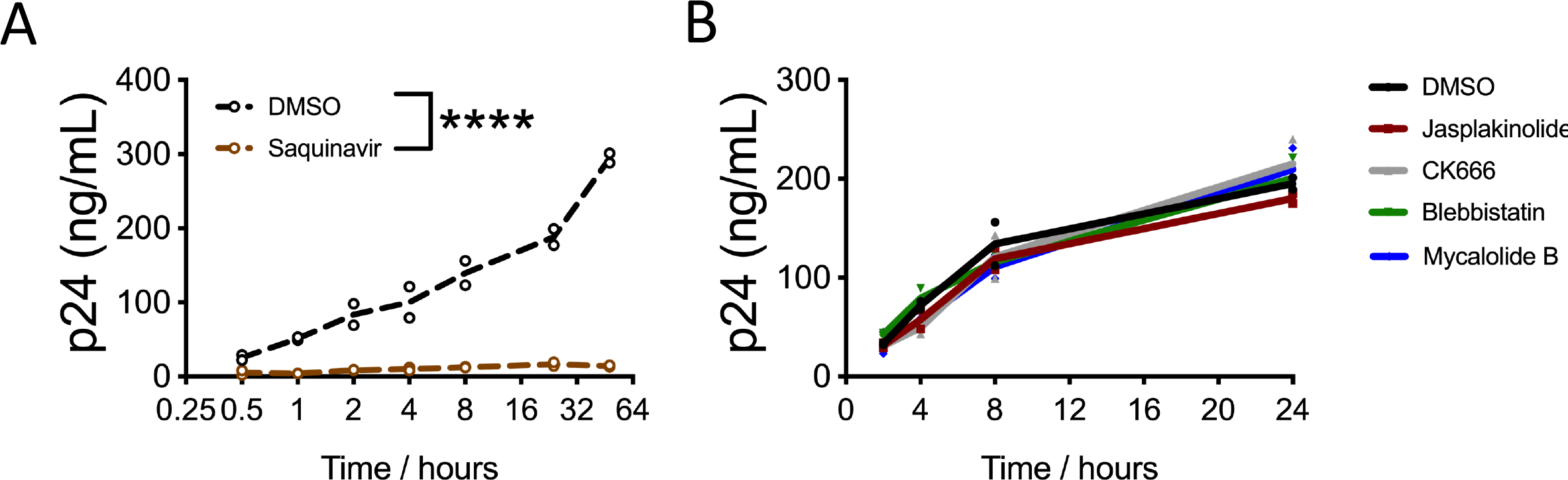

**Figure S3**. Validation of CBA assay and impact of actin modulation in HIV-1 release from HeLa cells

**A** – HEK 293FT cells were transfected with pNL-ΔENV-VSVG. After 16 hours media was replaced, and cell were treated with the viral protease inhibitor Saquinavir (10 µM) or vehicle (DMSO). Supernatants were recovered at the indicated time-points, filtered (0.45 µm) and p24 quantified via a custom-made CBA assay.

**B** – HeLa cells were infected with NL-ΔENV-VSVG (MOI=1.0). At 24 hours post-infection, culture media was replaced containing the indicated inhibitors or vehicle (DMSO). Aliquots of the supernatant were recovered at the indicated time-points, filtered (0.45 µm) and p24 quantified by CBA.

Jasplakinolide = 50 nM / CK666 = 10 µM / Blebbistatin = 5 µM / Mycalolide-B = 100 nM. Data from 3 independent experiments.

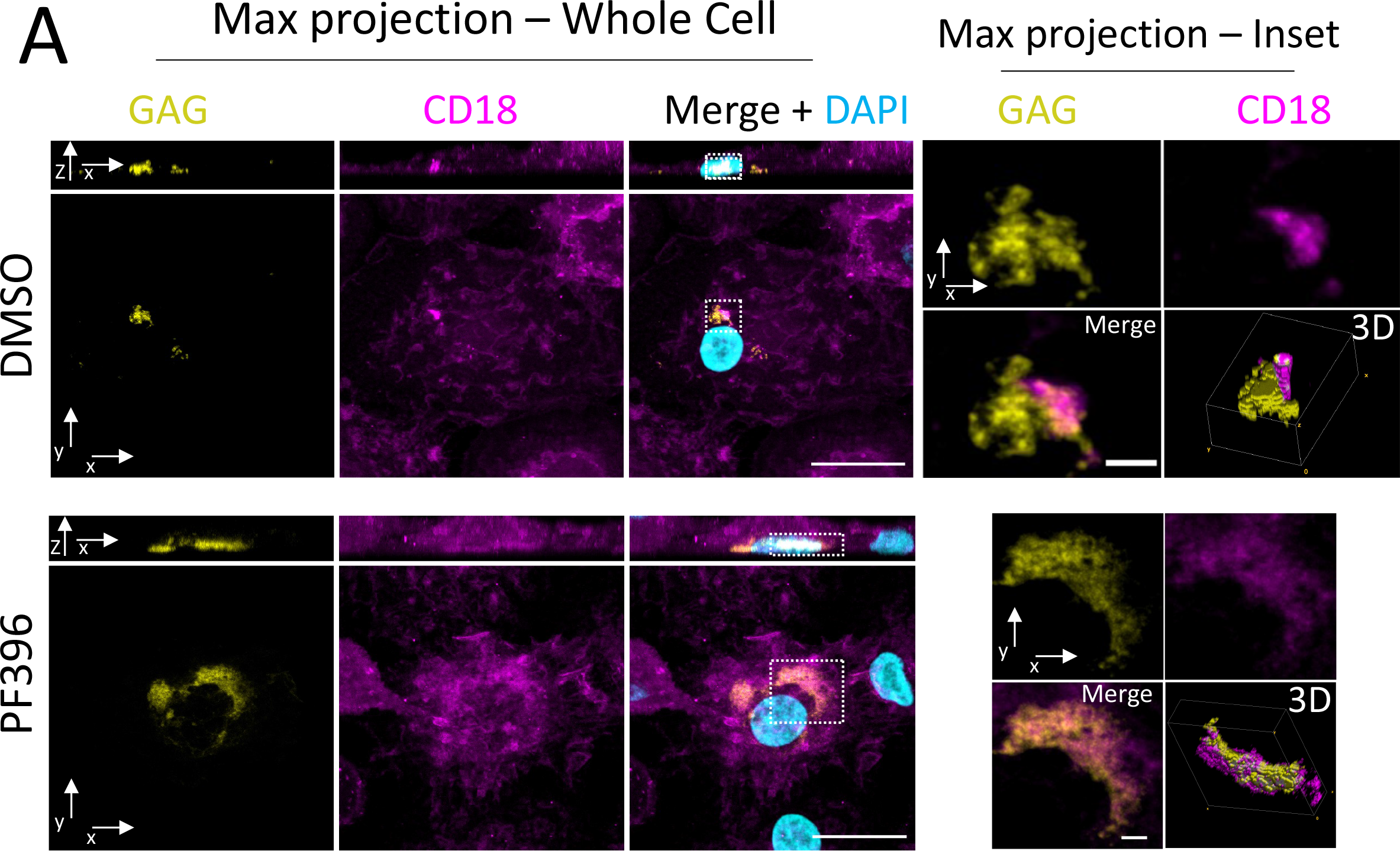

**Figure S4**. Altered distribution of CD18 in VCC after PF396 treatment

**A** – Confocal microscopy of MDMs infected with HIV-1-Gag-iGFP-ΔENV-VSVG (MOI=1.0) for 4 days and treated with PF396 (1µM) or vehicle (DMSO) for an additional 96 hours. Whole cell projections are shown on the right, along the z-axis (bottom) or the y axis (top). On the left, the areas enclosed in the dashed squares are zoomed in and shown as z-projections. Scale bar = 20 µm (left panels) or 2 µm (right panels).

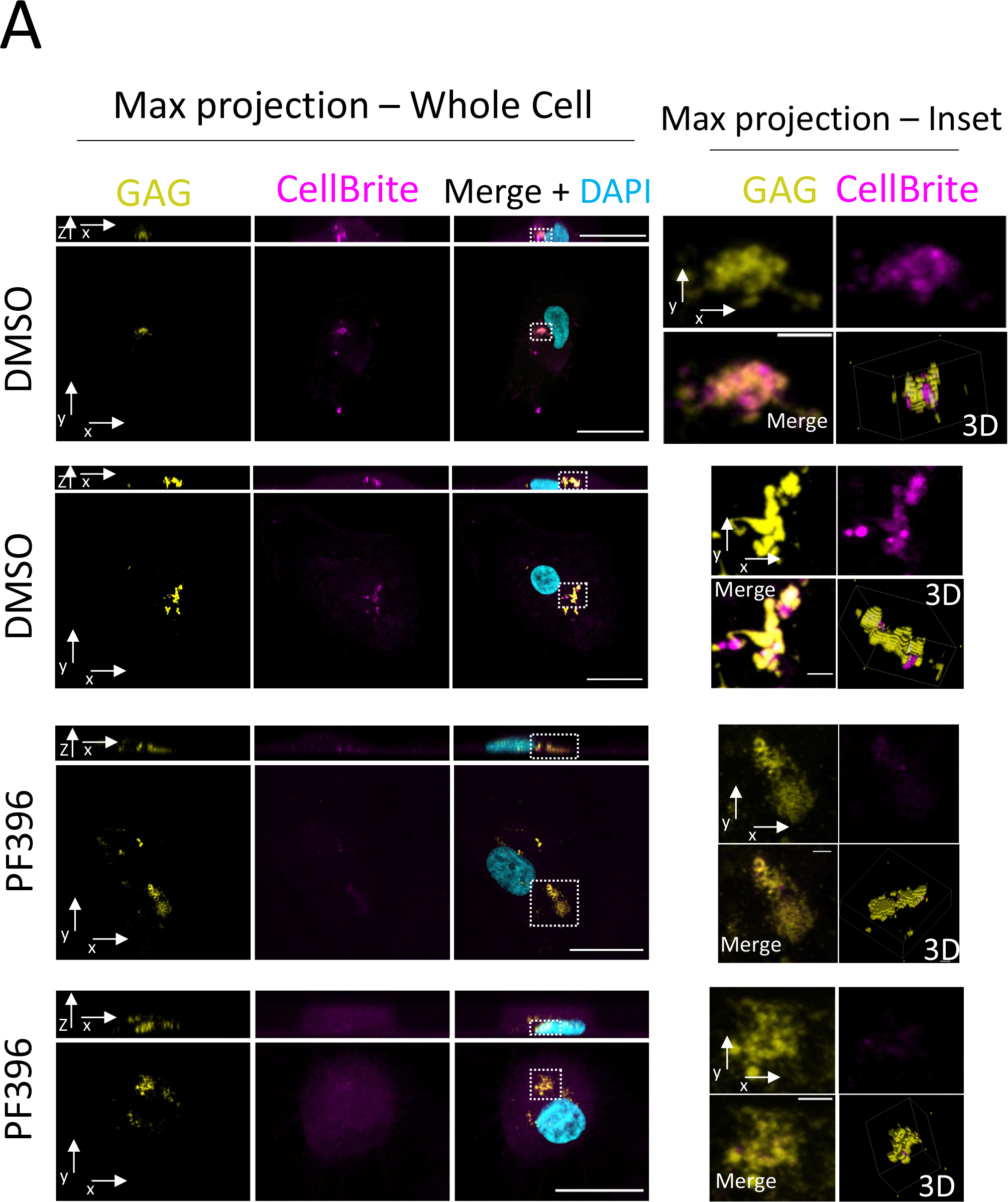

**Figure S5**. VCC in HIV-1-infected MDMs treated with PF396 lose connections with the membrane

**A** – Additional examples of CellBrite staining in MDMs infected with HIV-1-Gag-iGFP-ΔENV-VSVG and treated with PF396, as in Figure 5K.

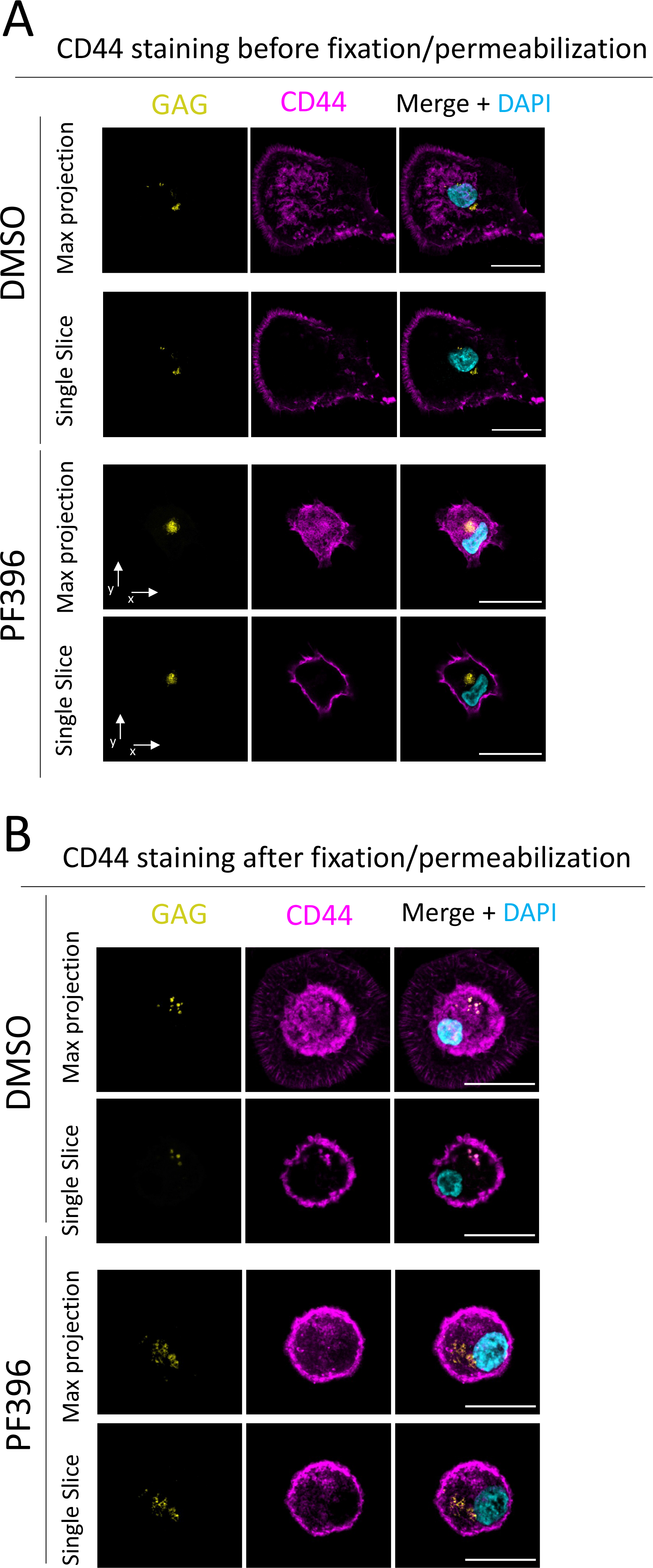

**Figure S6**. VCC remain inaccessible to antibodies delivered to the media after PF396 treatment.

**A** – Confocal microscopy of HIV-1-GAG-iGFP-ΔENV-VSVG-infected MDMs and treated with PF396 or DMSO. CD44 staining was performed before cell fixation and permeabilization.

**B** ~~–~~ Same as in A, except that CD44 staining was performed after cell fixation and permeabilization.

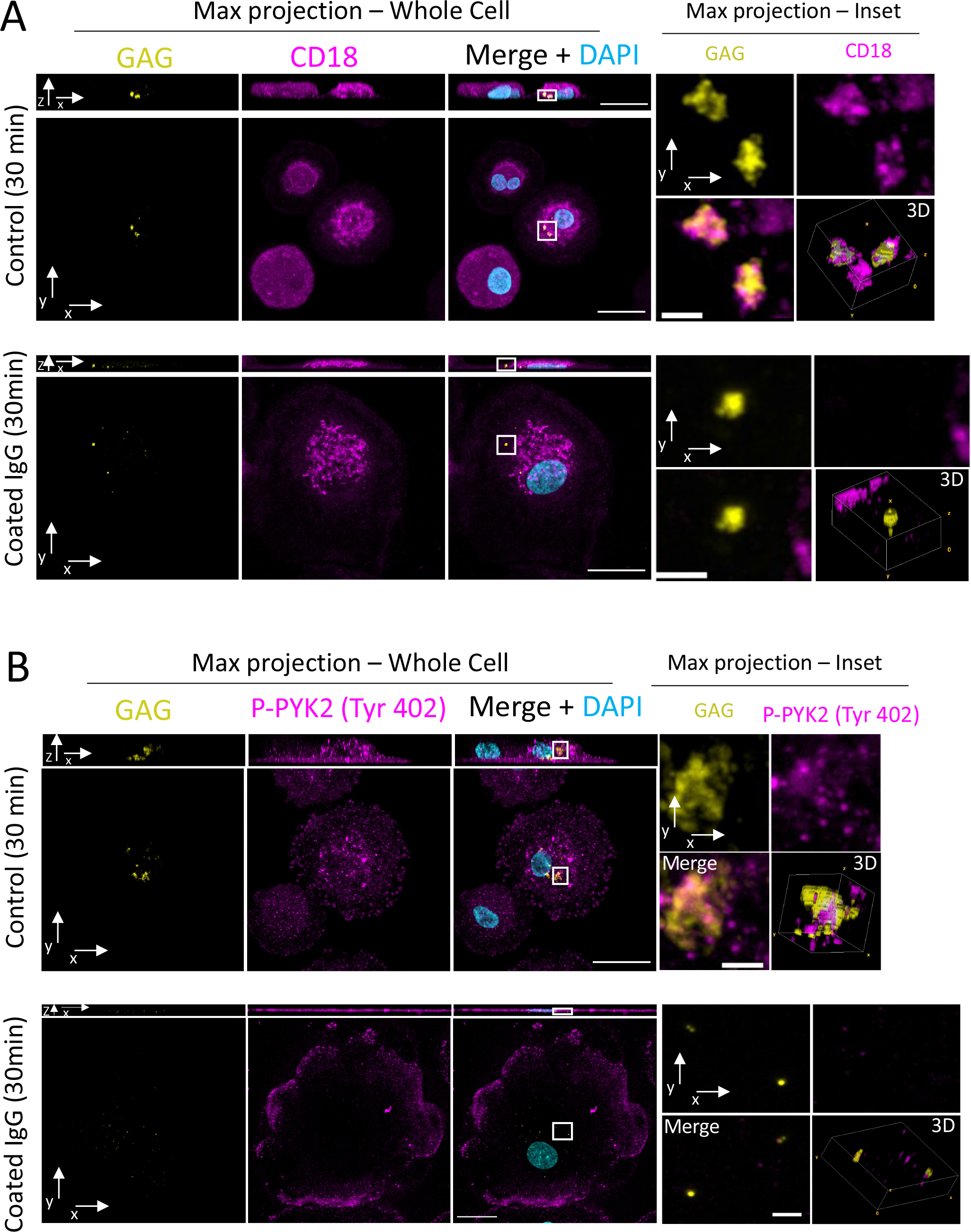

**Figure S7**. CD18 and p-PYK2 localization at the VCC before and after frustrated phagocytosis

**A-B** – Confocal microscopy of MDMs infected with HIV-1-Gag-iGFP-ΔENV-VSVG (MOI=1.0) for 3 days and seeded over control or human IgG-coated coverslips for 30 mins. Right panels - whole cell projections along the z-axis (bottom) or the y axis (top). Left, panels – magnification of the areas enclosed in the dashed squares as z-projections. Scale bar = 20 µm (left panels) or 2 µm (right panels).

**Video S1**: Dynamics of actin and VCC in HIV-1-infected macrophage

MDMs transduced with Lifeact-mCherry were infected with HIV-1-GAG-iGFP-ΔENV-VSVG (MOI=1.0). Time-lapse imaging was performed at day 6 after infection. Scale bar = 5 μm

**Video S2**. Live imaging of frustrated phagocytosis of HIV-1-infected macrophages.

Time-lapse imaging of MDMs infected with HIV-1-GAG-iGFP-ΔENV-VSVG for 3 days and seeded over human IgG-coats coverslips. Whole cell projections are shown for the top and middle panels, along the z-axis (middle) or the y axis (top). The bottom panel (transmission channel) is a single z-slice. Scale bar = 20 µm.

**Supplementary Table 1 – Antibodies employed in this study**

| **Antigen targeted** | **Application** | **Species** | **Dilution** | **Source** |
| --- | --- | --- | --- | --- |
| F-actin | Immunofluorescence | mouse | 1/100 | Abcam – ab205 |
| CD18 | Immunofluorescence | mouse | 1/100 | Abcam – ab657 |
| p-PYK2 (TYR402) | Immunofluorescence/ Western blot | rabbit | 1/100 (IF)  1/1000 (WB) | Abcam – ab4800 |
| CD44 | Immunofluorescence | rat | 1/100 | BD - 550538 |
| GAG | Western Blot | mouse | 1/2000 | NIH AIDS reagent resource - H183–H12–5C |
| GAPDH | Western blot | goat | 1/2000 | Abcam – ab157156 |
| PYK2 | Western Blot | rabbit | 1/1000 | Cell Signaling - 3292 |
| Clathrin heavy chain 1 | Western Blot | mouse | 1/1000 | Abcam – ab2731 |
| Anti-p24-FITC | CBA | mouse | 1/200 | Beckman Coulter / KC57 |
